# Ketogenic diet is protective during endotoxin-induced lung injury through the elevation of BHB

**DOI:** 10.64898/2026.09.20.753061

**Authors:** Robert Lwanga, Thu T. Tran, Jieun Kim, Katherine Figarella, Kevin Huang, Wanheng Zhang, Ragini Nair, Victor Guaregua, Sophia Phan, Heather Canales, Xuebo Chen, Synthea Horton, Yu A. An, Xiaoyi Yuan

## Abstract

Acute respiratory distress syndrome (ARDS) is marked by severe pulmonary edema and concomitant hypoxia, affecting hundreds of thousands of people a year, especially those in critical care conditions or suffering from septic shock. Previous studies have implicated that the ketogenic diet, a high-fat and low-carbohydrate diet, modulates inflammatory responses. However, the impact of the ketogenic diet on septic ARDS outcomes is unknown. Here, we demonstrated that mice on a ketogenic diet showed strikingly reduced lung injury and inflammation compared to those on a control diet during a murine model of endotoxin-induced lung injury, induced by intratracheal lipopolysaccharide (LPS) injection. Immune mass cytometry studies on lung tissue indicated that the ketogenic diet reduces immune cell infiltration. Treating mice with beta-hydroxybutyrate (BHB), the primary metabolite of ketogenesis, after the onset of ARDS reduced pulmonary edema and lung inflammation, as well as NF-kB activity, suggesting strong therapeutic potential. By multiplex analysis in bronchial alveolar lavage fluid, we observed that the ketogenic diet or BHB administration attenuates the chemotaxis and activation of immune cells. Altogether, our findings reveal that the ketogenic diet provides lung protection during endotoxin-induced lung injury through BHB.

## Main

Acute respiratory distress syndrome (ARDS) is caused by profound, uncontrolled lung inflammation with excessive immune cell infiltration and cytokine storm, eventually leading to severe pulmonary edema and poor oxygenation ^1,2^. ARDS impacts approximately 70 cases per 100,000 person-years in the US, and the mortality is high^3^. Sepsis is the most common cause of ARDS ^4^. So far, the treatment approaches for septic ARDS are mainly supportive care and controlling systemic infection, without effective treatment options to dampen excessive lung inflammation ^5^. Therefore, it is crucial to explore potential therapeutic approaches for septic ARDS.

Nutrition and metabolic support are essential components of management for critically ill patients, including those with ARDS. The ketogenic diet is characterized by a low-carbohydrate, high-fat diet with sufficient protein, which results in ketogenesis while maintaining caloric intake ^6^. Lately, the immunomodulatory impact of the ketogenic diet has been in the spotlight, including its involvement in several pathogenic conditions such as viral infection, neurological diseases, or cancer ^6-8^. Furthermore, recent studies imply that ketogenesis dampens excessive lung inflammation during *Pseudomonas aeruginosa* infection ^9^. Thus far, the safety of the ketogenic diet in the critically ill population has been supported by previous clinical studies^10^. However, the impact of the ketogenic diet on septic ARDS remains elusive.

Our study demonstrated the lung-protective effects of the ketogenic diet in murine models of endotoxin-induced lung injury, achieved via intratracheal injection of lipopolysaccharide (LPS). We also showcase the changes in the immune landscape modulated by the ketogenic diet via imaging mass cytometry. Furthermore, we showed the lung-protective effect of beta-hydroxybutyrate (BHB), the ketone body produced during ketogenesis, as a post-ARDS treatment. Finally, multiplex analysis identified several key inflammatory mediators modulated by the ketogenic diet and BHB.

### Ketogenic diet attenuates endotoxin-induced lung injury

Previous studies have suggested that a ketogenic diet may possess anti-inflammatory properties in several disease models. However, its impact on septic ARDS outcomes is currently unknown. Thus, we utilize a model of intratracheal injection of LPS to model septic ARDS in mice ^11^. To study the impact of ketogenic diet, we first confirmed that the duration of the diet successfully results in ketogenesis in mice. Indeed, 7 days of exposure to ketogenic diet in mice leads to significant weight loss and increases in blood levels of BHB, the primary metabolites from ketogenesis ^12^, compared to coco butter-based control diet, implicating conversion to ketogenesis (**Suppl Fig. 1a and b**). We then expose mice to endotoxin-induced lung injury model while keeping the mice on the same diet for three additional days following the initiation of lung injury (**Fig. 1a**). Mice under the ketogenic diet showed significant reduced weight loss following LPS exposure compared to the control diet, suggesting improved outcomes (**Fig. 1b**). Ketogenic diet also reduces pulmonary edema marked by reduced total protein and albumin levels in the bronchoalveolar lavage (BAL) fluid (**Fig. 1c and d**). There is also a significant decrease in inflammatory cell infiltration into the airway in the ketogenic group, as both total BAL leukocytes and neutrophils are lower in the ketogenic group compared to the control diet (**Fig. 1e and f**). The overall attenuation of lung injury and inflammation in the ketogenic diet is also associated with improvement in histological evidence of lung injury, as mice under the ketogenic diet show lower alveolar overdistension, atelectasis, cellularity, and edema (**Fig. 1g to i**). Thus, the ketogenic diet provides lung protection during murine endotoxin-induced lung injury.

**Figure 1:**
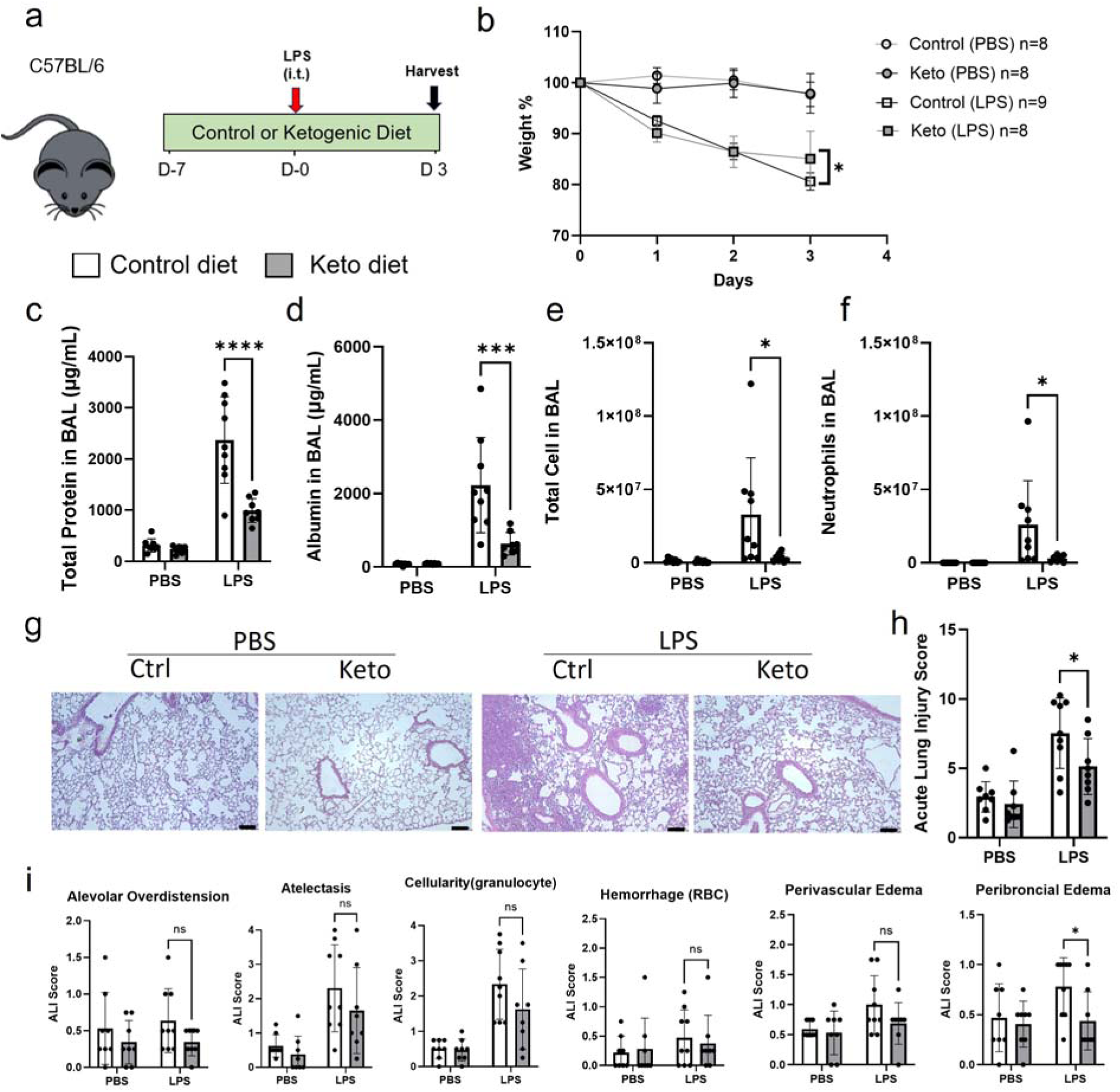
Ketogenic diet attenuates endotoxin induced lung injury. **a.** Experimental schematic. C57BL/6 mice were established on a 7-day ketogenic (keto) diet, followed by a 3-day LPS-induced ARDS. Mice were harvested after 3 days post-LPS for blood serum, bronchoalveolar lavage fluid (BALF), and lungs. **b.** Weight loss curve of mice on a 3-day LPS-induced ARDS with four groups: Mice on a keto diet that were injected with and without LPS, and control diet mice that were injected with and without LPS(n= 8 in the Control diet PBS group, n= 8 in the Ketogenic diet PBS group, n=9 in the Control diet LPS group, n= 8 in the Ketogenic diet LPS group)**. c.** Total protein level extracted from BALF using BCA protein assay.(n=8-9 per group, two-way ANOVA with Šídák’s test. **d.** Albumin levels from BALF via ELISA. (n=8-9 per group, two-way ANOVA with Šídák’s test.) **e.** Total cell count of BALF. (n=8-9 per group, two-way ANOVA with Šídák’s test.) **f.** Neutrophil cell count of BALF cytospin (n=8-9 per group, two-way ANOVA with Šídák’s test.) **g.** 10x magnification of images of H&E staining of the lungs. Scale bar is 100 μm. **h.** ARDS score of the H&E-stained images. (n=8-9 per group, two-way ANOVA with Šídák’s test.) **i.** Subcategories that were used for the ARDS scoring of the H&E-stained slides from mice on a 3-day LPS-induced ARDS that include: alveolar overdistension, atelectasis, cellularity, hemorrhage, perivascular edema, and peribronchial edema. (n=8-9 per group, two-way ANOVA with Šídák’s test.) All data are represented as mean ± SD; *P-value < .05; ns p-value >0.05.

### Ketogenic diet modulates immune cell infiltration

So far, our study implicated that the ketogenic diet mediated lung protection during endotoxin-induced lung injury. Next, we further dissect the immune cell infiltration into the lung via imaging mass cytometry ^13^. Lungs were embedded in paraffin and labeled with 21 heavy metal-tagged antibodies (**Suppl Fig. 2a**). Firstly, we created a 2D map of specific cell populations based on the molecular targets of the antibodies (**Fig. 2a**). We then further investigated changes in particular cell types comparing the ketogenic diet and control diet under PBS or LPS exposure conditions. We have observed a significant reduction of several immune cell populations in the ketogenic diet group during endotoxin-induced lung injury, including Ly6G+, MHCII+, F480+, F480+/Arg1+, F480+/CD206+, and CD4+ Foxp3+ cell populations (**Fig. 2b**). The reduced Ly6G+ cells are consistent with the reduction of neutrophils observed in the ketogenic diet group during endotoxin-induced lung injury. Interestingly, regulatory T cells (CD4^+^/Foxp3^+^) are also reduced in the ketogenic diet group (**Fig. 2b**), which is unexpected since these cells are defined as immune regulatory cells, and the ketogenic diet group experienced less lung inflammation. However, several previous reports suggested that the population of regulators increases with the degree of inflammation ^14-16^. Thus, the reduction of Treg population could directly correlate with the lessened lung inflammation in the ketogenic diet group. Interestingly, the inhibition of immune cell infiltration is not a global impact, as several immune cell populations are enhanced in the ketogenic diet group. For example, CD163^+^ populations are significantly higher in the ketogenic diet group (**Fig. 2b**). CD163 is a macrophage scavenger receptor that is highly expressed on alternatively activated M2 macrophages ^17^. Finally, several immune cell population remains unchanged comparing different diet during endotoxin induced lung injury, including B cells (B220^+^), pan T cells (CD3^+^) natural killer cells (NK^+^), and dendritic cells (CD11c^+^) (**Suppl Fig. 2b**). There is also no differences in Ki67 ^18^ or alpha-SMA ^19^ expression, indicating that ketogenic diet does not alter cell replication or the fibrogenic activity (**Suppl Fig. 2b**). Thus, ketogenic diet inhibits infiltration of inflammatory cells, implicating potential impacts on immune cell chemotaxis.

**Figure 2:**
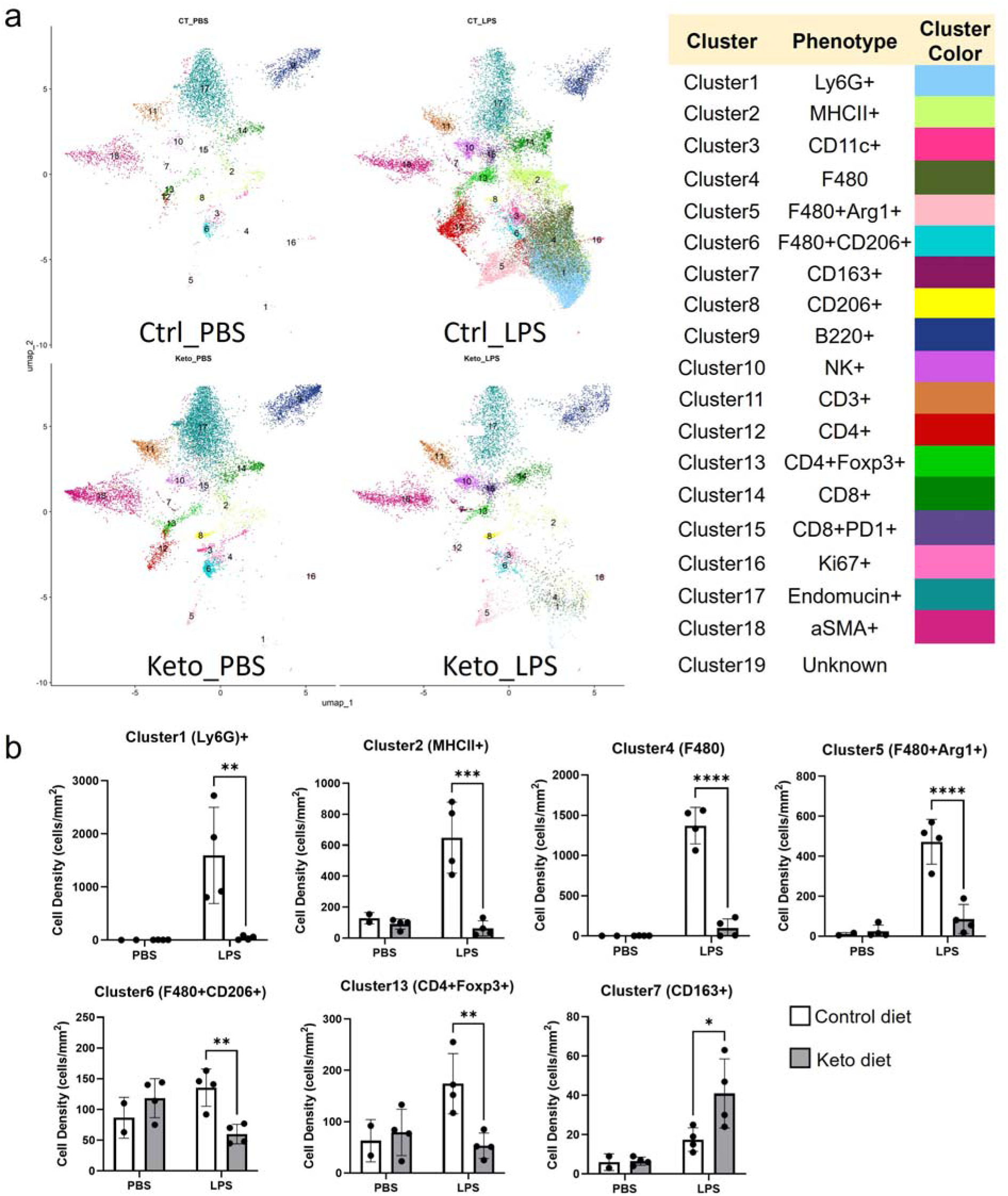
Ketogenic diet modulates immune cell infiltration. **a.** Uniform Manifold Approximation and Projection (UMAP) of mice on a 3-day LPS-induced ARDS with four groups: Mice on a ketogenic (keto) diet that were injected with or without LPS, and control diet mice that were injected with or without LPS**. b.** Significantly different immune cell populations based on the UMAP of mice on a 3-day LPS-induced ARDS with four groups: Mice on a keto diet that were injected with or without LPS, and control diet mice that were injected with or without LPS. (n=2 in the Control diet PBS group, n=4 in the Keto diet PBS group, n=4 in the Control diet LPS group, n=4 in the Keto diet LPS group. Two-way ANOVA with Šídák’s test.) All data are represented as mean ± SD; *P-value < 0.05

### Post-ARDS beta-hydroxybutyrate (BHB) treatment alleviates lung injury and inflammation

BHB is one of the primary metabolites during ketogenesis. Several studies have suggested that BHB is essential for the anti-inflammatory effect mediated by the ketogenic diet ^20-22^. Thus, we investigated whether direct administration of BHB could recapitulate the lung protection observed in a ketogenic diet during endotoxin-induced lung injury. Since the half-life of BHB is relatively short ^23^, mice were treated with intraperitoneal injection of 2521.8 mg/kg of BHB twice daily. The injection successfully elevated BHB levels in the serum 2 hours after the injection (**Suppl Fig. 3**). Starting one day after the initiation of endotoxin-induced lung injury, mice received BHB injection twice per day until the final harvesting at day 4 post-LPS injection (**Fig. 3a**). We observed less weight loss in BHB treated mice compared to control vehicle (PBS) treated group on day 4 post LPS injection (**Fig. 3b**). Moreover, BHB-treated mice have a significant reduction of pulmonary edema, marked by lower levels of albumin in the BAL (**Fig. 3c**). Furthermore, analysis of the composition of BAL leukocytes indicated an overall dampening of immune cell infiltration with a significant reduction of neutrophils, consistent with those maintained under the ketogenic diet (**Fig. 3d-e**). Furthermore, the histological analysis of lung tissue revealed that the BHB treatment group had reduced lung injury compared to the PBS control (**Fig. 3f-h**). The profound protection is only observed in mice receiving 3 days of BHB treatment, as mice receiving 2 days of BHB treatment and harvested 72 hours post LPS injection showed similar lung injury and inflammation compared to the vehicle control group (**Suppl Fig. 4**). Overall, our results demonstrate that BHB, a metabolite produced during ketogenesis, plays a crucial role in alleviating lung injury and inflammation mediated by the ketogenic diet during endotoxin-induced lung injury.

**Figure 3:**
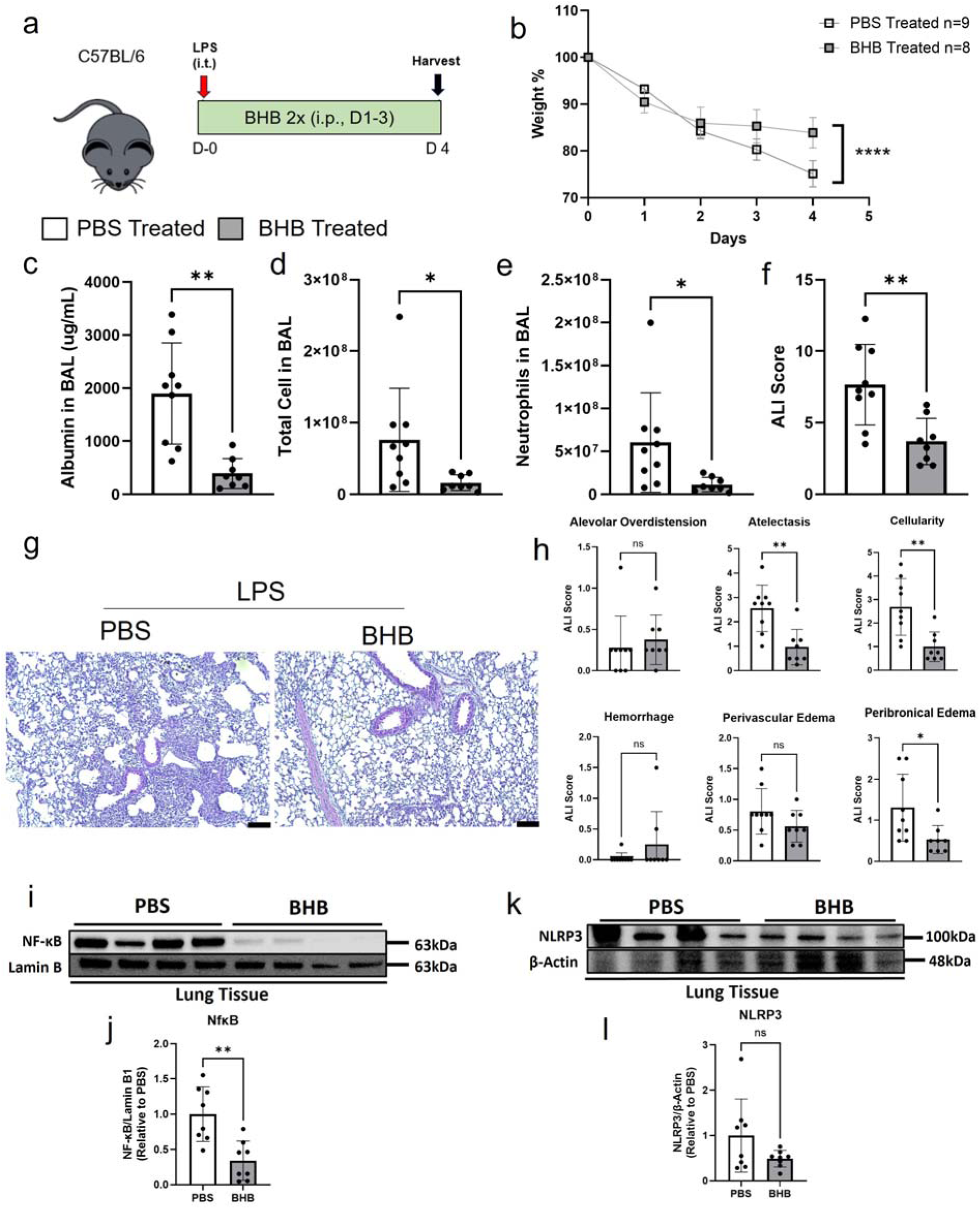
Beta-hydroxybutyrate (BHB) post-ARDS treatment attenuates lung injury and inflammation. **a.** An experimental layout. C57BL/6 mice had LPS instilled intratracheally on day 0 and followed for 4 days. Mice were injected with BHB twice a day for 3 days starting on day 1. Mice were harvested for their blood serum, BAL fluid, and lungs. **b.** Weight loss curve of mice on a 4-day LPS-induced ARDS with two groups: Mice injected with BHB twice a day, and mice that were injected with PBS twice a day. (n=9 in the PBS group, n=8 in the BHB-treated group) **c.** Albumin levels extracted from BALF from mice on a 4-day LPS-induced ARDS. (n=8-9 per group, Mann-Whitney U test.) **d.** Total cell count of BALF from mice on a 4-day LPS-induced ARDS. (n=8-9 per group, Mann-Whitney U test.) **e.** Neutrophil cell count of BALF cytospin from mice on a 4-day LPS-induced ARDS. (n=8-9 per group, Mann-Whitney U test.) **f.** ARDS score of the H&E-stained images. (n=8-9 per group, unpaired t-test with Welch’s correction.) **g.** 10x magnification of images of H&E staining of the lungs. **h.** Subcategories that were used for the ARDS scoring of the H&E-stained slides include: alveolar overdistension, atelectasis, cellularity, hemorrhage, perivascular edema, and peribranchial edema. (n=8-9 per group, Mann-Whitney U test or unpaired t-test with Welch’s correction.) **i.** and **j.** Image and densitometry quantification for Western blot for NF-κB protein expression in lung tissue isolated from BHB-treated mice compared to PBS-treated mice after 4-day LPS-induced ARDS. (n=8 in the PBS-treated group, n=8 in the BHB-treated group. Unpaired t-test with Welch’s correction) **k.** and **l.** Image and densitometry quantification for Western blot for NLRP3 protein expression in lung tissue isolated from BHB-treated mice compared to PBS-treated mice after 4-day LPS-induced ARDS. (n=8 in the PBS-treated group, n=8 in the BHB-treated group. Unpaired t-test with Welch’s correction.) All data are represented as mean ± SD; *P-value < 0.05; ns p-value >0.05.

So far, our study has found that BHB and the ketogenic diet can alleviate inflammation and lung injury; however, the specific molecular mechanisms remain unclear. Thus, we conducted studies investigating the impact of BHB treatment on key signaling pathways, including the NF-kB and NLRP3 inflammasome. Consistent with the reduction of lung inflammation, NF-kB protein level is significantly reduced in the lung tissue from BHB-treated mice during LPS-induced lung injury (**Figure 3i, j**). To investigate whether canonical NF-kB activation is the main player for the differences in NF-kB protein levels between PBS and BHB-treated groups, we performed western blotting of phosphorylated IκBα ^24^. Surprisingly, there is no difference in the phosphorylation of IκBα (**Suppl Fig. 5**), suggesting that the acute canonical NF-kB activation is not the main mechanism. BHB could potentially inhibit NF-kB activation through non-canonical pathways or by disrupting the self-sustained positive feedback loop of NF-kB ^25^. Previous studies also indicated that BHB exerts anti-inflammatory effects by dampening NLRP3 activity ^20,26^. However, we observed no significant reduction of NLRP3 levels in the BHB-treated mice (**Figure 3k, l**). Overall, this study suggested that NF-kB signaling might underlie BHB-mediated lung protection during endotoxin-induced lung injury. However, the detailed molecular mechanisms require further investigation.

### Ketogenic diet and BHB post-ARDS treatment reduce inflammatory mediators

Next, we conducted a multiplex assay^27^ on bronchial alveolar lavage fluid to quantify changes in chemokines and cytokines under different dietary conditions or BHB treatment groups during endotoxin-induced lung injury to investigate the underlying mechanism for lung protection. The ketogenic diet reduced the levels of eotaxin, granulocyte-macrophage colony-stimulating factor (GM-CSF), LIF, and MCP-1 compared to the control diet (**Fig. 4a**). These results suggest that the ketogenic diet reduces the chemotaxis and differentiation of myeloid cells, which is in line with our immune mass cytometry analysis. Next, we investigate the impact of BHB post-ARDS treatment on cytokine/chemokine levels in bronchial alveolar lavage fluid at day 4 post-endotoxin-induced lung injury. Consistently, BHB treatment also reduced eotaxin, GM-CSF, and LIF (**Fig. 4b**), suggesting a similar pathway that is suppressed by BHB. Interestingly, BHB treatment dampened several other inflammatory mediators that were unaltered by ketogenic diet pretreatment, including MIP-1a, MIP-1b, RANTES, IL-1a, IL-1b, IP-10, M-CSF, IFNg, and TNFa (**Fig. 4b**). Cytokines that are unaltered are included in **Suppl Fig. 6 and 7**. The increased potency and broader anti-inflammatory effects of BHB treatment suggest it may be a superior treatment option, especially after the onset of ARDS. Taken together, based on a multiplex assay of the BAL fluid, both the pre-ARDS ketogenic diet treatment and the post-ARDS BHB treatment attenuate immune cell chemotaxis and activation.

**Figure 4:**
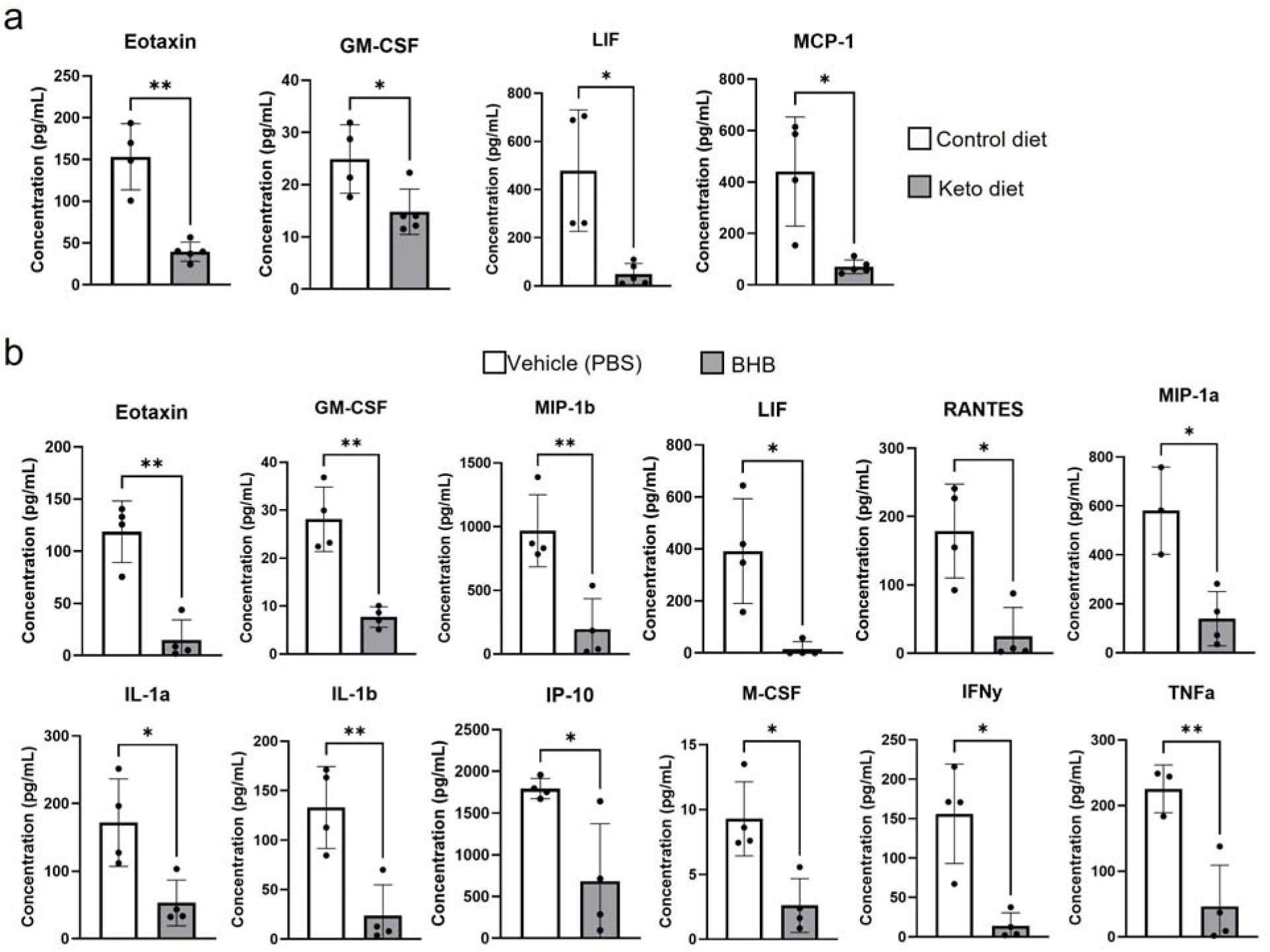
Ketogenic Diet or BHB post-ARDS treatment reduces inflammatory mediators during endotoxin-induced lung injury. **a.** List of significantly different cytokines and chemokines from the Luminex mouse cytokine immunoassay in bronchoalveolar lavage fluid (BALF) from ketogenic (keto) diet mice compared to control mice after 3-day LPS-induced ARDS. (n=4 in the Control diet LPS group, n=5 in the Ketogenic diet LPS group. Unpaired t-test with Welch’s correction.) **b.** List of significantly different cytokines and chemokines from the Luminex mouse cytokine immunoassay in bronchoalveolar lavage fluid (BALF) from BHB-treated mice compared to PBS-treated mice after 4-day LPS-induced ARDS. (n=4 in the PBS-treated group, n=4 in the BHB-treated group. Unpaired t-test with Welch’s correction.) All data are represented as mean ± SD; *P-value < 0.05, **P-value < 0.01.

## Discussion

Septic ARDS has high mortality and morbidity in the critically ill population, and therapeutic treatment is still lacking. Modification of nutrition is a feasible approach for managing septic ARDS in the ICU. However, the impact of the ketogenic diet on septic ARDS remains unexplored. Our study demonstrated that a ketogenic diet offered lung protection in a murine model of septic ARDS, resulting in a significant reduction in lung injury and inflammation. Additional imaging mass cytometry analyses implicated significant attenuation of immune cell chemotaxis in ketogenic diet-treated groups. Interestingly, the ketone body BHB recapitulates the lung protection observed in the ketogenic diet condition when administered after the initiation of ARDS. Finally, multiplex assay suggested that both the ketogenic diet and BHB post-ARDS treatment attenuate immune cell activation and chemotaxis.

Our study suggests that the ketogenic diet or BHB post-ARDS treatment provides lung protection during a murine model of endotoxin-induced lung injury. These results are consistent with the immune-modulatory effects of ketogenesis established in previous publications ^7,21,28^. However, the detailed signaling pathways remain to be further explored. Our studies have suggested that BHB treatment is associated with reduced NF-kB activity. However, the detailed molecular mechanism of BHB-mediated NF-kB inhibition in the context of endotoxin-induced lung injury remains unclear. And identifying the key cellular bases of protection is one of the prerequisites to pinpointing the detailed molecular mechanism, which also requires further investigations. Furthermore, whether this response is related to BHB receptors remains unclear. Indeed, previous studies have suggested that dimethyl fumarate (a GPR109A agonist) may reduce LPS-induced lung injury ^29^. Thus, it is reasonable to hypothesize that BHB might mediate lung protection during endotoxin-induced lung injury through directly binding its receptors. However, additional studies are required to test this hypothesis, and at this stage, its involvement is purely speculative, rather than an inferred mechanism. Moreover, hypoxia inducible factor (HIF) stabilizes during ARDS via various mechanisms, including low oxygen levels or succinate accumulation ^30^. HIF stabilization is crucial for lung protection during ARDS ^2,27,31,32^. As previous studies indicated that a ketogenic diet could increase the levels of succinate, it could potentially lead to HIF stabilization and then lung protection ^33^. However, further studies are required to elucidate the signaling pathways crucial for the ketogenic diet and BHB-mediated lung protection.

The lung protection afforded by the ketogenic diet in septic ARDS could result from pulmonary or extrapulmonary mechanisms. As a dietary modification, the metabolic changes induced by the ketogenic diet are likely to be systemic, affecting all cells in the body. Since metabolic optimization is crucial for alveolar epithelial or immune cell functions ^34,35^. The lung protection observed in the ketogenic diet likely stems from the different metabolite profiles that occur during ketogenesis or the metabolic reprogramming of lung cells. On the other hand, since the liver is one of the key organs for metabolism ^36^, it could be responsible for the changes in lung metabolic microenvironment during the ketogenic diet, and contribute to the subsequent changes in immune responses in the lung during endotoxin-induced lung injury. The fact that BHB recapitulates the lung protection observed in the ketogenic diet suggests that metabolites from either lung-resident cells or other organs are crucial in dampening pulmonary injury during septic ARDS. Finally, we cannot rule out the potential involvement of the microbiome in the gastrointestinal tract, as it could alter the profile of metabolites circulating systemically under a ketogenic diet, contributing to changes in the lung metabolic microenvironment. Indeed, a recent study suggested that a ketogenic diet alleviates septic lung injury via the microbial gut-lung axis ^37^. Further studies are needed to elucidate those important mechanisms.

Our study has its limitations and challenges that will need to be addressed for further translation into the clinic. First, some of the n numbers in the IMC analysis are small with n=2. Thus, additional confirmatory studies are needed to confirm the changes in several key immune cell populations. The reduction in Tregs (CD4^+^ Foxp3^+^ cells) during a ketogenic diet creates a potential paradox with established mechanisms, as Tregs typically exert anti-inflammatory functions. Additional studies are needed to validate Treg functionality (e.g., IL-10 secretion assays) or employ Treg-depletion models to clarify their role. Furthermore, our findings will need to be replicated in other septic ARDS models, including the cecal ligation and puncture model or the fecal slurry injection model. These two models involve systemic live bacterial infection, and the fact that ketogenesis is protective against Pseudomonas pulmonary infection suggests that the ketogenic diet may have a similar protective effect observed in endotoxin-induced lung injury. Our study focuses on the pretreatment of mice on the ketogenic diet prior to the initiation of endotoxin-induced lung injury, which could be translated into patients who are at high risk of developing ARDS during elective surgical procedures. However, additional studies are needed to fine-tune the timeline of the ketogenic diet for its use as a treatment for ARDS, including identifying the shortest diet maintenance time required for the lung-protective effect. We demonstrate that BHB treatment initiated after ARDS onset provides lung protection. The BHB dosage used in our study is derived from previous literature. However, whether our current treatment strategy is optimal in terms of dosage and timing requires further investigation. Given BHB’s short half-life, additional studies should also focus on pharmacologic modulators of the BHB receptor. Thus, treatment approaches based on BHB could significantly facilitate the translation of our findings, as they are readily available ^22^.

Taken together, we have identified the lung protective effect of the ketogenic diet during murine endotoxin-induced lung injury with a significant attenuation of immune cell chemotaxis. Our work further identified ketone body BHB as one of the key metabolites during ketogenesis responsible for the anti-inflammatory effects mediated by the ketogenic diet. This paved the way for further investigations into the cellular and molecular mechanisms mediated by the ketogenic diet.

## DECLARATION OF GENERATIVE AI AND AI-ASSISTED TECHNOLOGIES IN THE WRITING PROCESS

None to declare.

## CREDIT AUTHORSHIP CONTRIBUTION STATEMENT

- Conceptualization: XY
- Data curation: RL, JK
- Formal analysis: RL
- Funding acquisition: XY
- Investigation: KH, WZ, VG, XC, SH, SP, HC
- Methodology: RL, YAA
- Project administration: RN
- Resources: XC
- Supervision: XY
- Validation: KF, TTT
- Visualization: RL
- Writing – original draft: XY, RL
- Writing – review and editing: JK, KF, TTT, KH, WZ, YAA

## DATA AVAILABILITY

Data is included in the attached Excel sheet.

## METHODS

### Mice

C57BL/6J (Wildtype, female and male) mice aged 8 to 12 weeks were used for experiments. Mouse procedures were approved by the Institutional Animal Care and Use Committee at the University of Texas Health Science Center at Houston (AWC-23-0058). Mice were purchased from Jackson Laboratories. Once purchased, mice were housed and bred in a pathogen-free environment at the Center for Laboratory Animal Medicine and Care at the University of Texas Health Science Center at Houston ^38^.

### Ketogenic diet and LPS Model

A total of 33 mice were evenly split into two groups: Ketogenic and Control. The ketogenic group was fed a mostly cocoa butter-based ketogenic diet (D10070801, Research Diets, Inc., New Brunswick, USA) for 7 days, while the control group was fed a control diet provided by the same company (D10070802, Research Diets, Inc., New Brunswick, USA) for 7 days ^39,40^. Both groups’ weight was monitored every day. To establish the lung injury model, mice were intratracheally injected with 3.5 mg/kg lipopolysaccharide (LPS) (Millipore Sigma, L4391) ^38^. Mice were anesthetized with isoflurane. Three mice were randomly selected from both groups to receive the LPS injection, while the remaining two from each group received the same volume of PBS. The weights and wellness of the mice were monitored post-injection for 3 days, and then they were euthanized. A tracheostomy was performed after euthanasia, in which the bronchoalveolar lavage fluid (BALF) was collected by instilling and withdrawing 500 μL of PBS directly into the trachea of the mouse 3 times, and then centrifuged for 12 minutes at 1,200 rpm. Then the BALF was separated from the BAL pellet, and both were stored at -80 °C. Blood was collected from the heart and then centrifuged for 12 minutes at 1,200 rpm at 4 °C, and then the supernatant serum was collected and stored at - 80 °C. The right lungs were cut out of the mouse and then snap frozen via liquid nitrogen and then stored at -80 °C. The left lung was filled with formalin and stored in formalin.

### BAL differential

BAL was used to acquire a cell count using LUNA-II Automated Cell Counter (Logos Biosystems) and was used to create cytospin slides using Rotofix 32 A (Hettich) and then was stained with HEMA 3. Stained slides were visualized using a Leica Microscope, and a total of 200 neutrophils and macrophages were counted ^41^. The Neutrophil: Macrophage ratio was used with their respective cell count to generate an absolute cell count of neutrophils.

### Total Protein & Albumin Quantification

Total protein concentration in mouse BALF was measured using the BCA kit (Thermo scientific, 23225). Albumin concentration in mouse BALF was measured using enzyme-linked immunosorbent assays (Bethyl Laboratories) ^38^.

### Lung Histology

After harvest, the left lung of all mice was kept submerged in 10% formaldehyde (Epredia, 5701) for 24 – 48 hours and was then processed using a Leica TP1029 Automatic Benchtop Tissue Processor (Leica Camera, Wetzlar, Germany). After processing, the Lung tissue was cut into 5-micrometer sections and then stained with hematoxylin (Epredia 5701) and eosin (Epredia 71311) (H&E). Images of the stained sections were captured using a Leica Microscope (DMC 2500LED). Stained sections were scored for levels of acute lung injury by two scorers who were blinded to the experimental information (no injury = 0; injury to 20% of lung = 1; injury to 40% of lung = 2; injury to 60% of lung = 3; injury to 80% of lung = 4; injury to 100% of lung = 5). Each section was scored on 6 different categories: Alveolar Overdistension, Atelectasis, Cellularity, Hemorrhage, Perivascular Edema, and Peribronchial Edema. On a scale of 0-5 for each category, with 0 being no damage and 5 being complete injury to the entire lung per category. ^38^.

### Imaging Mass Cytometry (IMC) Analysis

Imaging Mass Cytometry (IMC) analysis was performed at the ImmunoMonitoring Core at the Houston Methodist Neal Cancer Center, Houston Methodist Research Institute. Antibodies were conjugated with metals according to the manufacturer’s protocol (Standard BioTools) as previously described^42^. Sample sections were baked at 60 °C overnight, and then deparaffinized in xylene and rehydrated in a serial gradient of alcohol for 10 min each. Epitope retrieval was done in a water bath at 95 °C in Tris-Tween 20 buffer, pH 9, for 20 min. Sections were blocked with 3% BSA in tris-buffered saline, and stained overnight at 4 °C with selected markers and counterstained with Cell-ID Intercalator (Standard BioTools). Slides were air-dried and ablated with Hyperion (Standard BioTools) for data acquisition. The IMC data are preprocessed and checked for tissue integrity, staining quality, and signal intensities prior to analysis.

For every region of interest (ROI), the tiff files are processed with *Steinbock* and the single cells are segmented using *Mesmer*, based on DNA staining (Ir191) and other cell surface markers ^43,44^. Following cell segmentation, the *Histology Topography Cytometry Analysis Toolbox (HistoCAT)*, are used for extracting single cell protein expression matrix ^45^. Data from all samples are consolidated in R scripts for downstream analysis. The normalized and standardized data are used to perform dimension reduction and unsupervised clustering in Seurat ^46^. Cell clusters are annotated based on marker expressions. Cell densities of each cell type are calculated by normalizing cell counts by areas (or mm^2^) of ROIs.

### Multiplex assay

Mouse BALF was used for measuring cytokine & chemokine profiling using a multiplex assay (MILLIPLEX® Mouse Cytokine/Chemokine Magnetic Bead Panel - Premixed 32 Plex-MCYTMAG70PMX32, Millipore Sigma) according to the manufacturer’s suggested protocol.

### BHB post-ARDS treatment

A total of 17 mice were split into two groups: BHB (DL-β-Hydroxybutyric acid sodium salt purchased from Millipore Sigma: H6501) and PBS. To establish the lung injury model, all mice were intratracheally injected with 3.5 mg/kg LPS. After the injury model was established, both groups were intraperitoneally injected with 2521.8 mg/kg of their respective solutions twice a day for a total of two or three days ^47^. Mice were anesthetized with isoflurane. The weights and wellness of the mice were monitored post-injection for four days, and then they were euthanized. A tracheostomy was performed after euthanasia, in which the bronchoalveolar lavage fluid (BALF) was collected by instilling and withdrawing 500 μL of PBS directly into the trachea of the mouse 3 times, and then centrifuged for 12 minutes at 1,200 rpm. Then the BALF was separated from the BAL pellet, and both were stored at -80 °C. Blood was collected from the heart and then centrifuged for 12 minutes at 1,200 rpm at 4 °C, and then the supernatant serum was collected and stored at -80 °C. The right lungs were cut out of the mouse and then snap frozen via liquid nitrogen and then stored at -80 °C. The left lung was filled with formalin and stored in formalin.

### Western blotting

Protein was extracted using NE-PER Nuclear and Cytoplpasmic Extractions Reagents (Thermo Fisher, 78835). Protein was quantified using Bradford Reagent (Bio-Rad, 5000205). After electrophoresis, membranes were blocked for 1 hour at room temperature in EveryBlot Blocking Buffer (Bio-Rad, 12010020). Membranes were incubated in primary antibodies at 4°C overnight with NF-kappaB p65 (6956), Lamin B1 (12586), NLRP3 (15101), IκBα (4814), Phospho-IκBα (2859), and beta-Actin (4970), all purchased from Cell Signaling Technology. Horseradish peroxidase-conjugated secondary antibodies (anti-rabbit IgG, HRP-linked antibody, 7074 or anti-mouse IgG, HRP-linked antibody, 7076, Cell Signaling Technology) were used at 1:2500 dilution. Membranes were imaged on a Bio-Rad ChemiDoc Touch Imaging System. ImageJ software (National Institutes of Health) was used for protein quantification.

### Statistical Analysis

Data were analyzed by using GraphPad Prism (version 10, San Diego, CA). For data only comparing two groups, normality tests are performed. If data are normally distributed and the equal-variance assumption is not met, an unpaired t-test with Welch’s correction was used. If the data were not normally distributed, the Mann-Whitney test was used. *P*-values less than .05 were considered statistically significant. For data comparing more than two groups, a two-way ANOVA was performed, and Šídák’s test was used for multiple comparisons. *P*-values less than .05 were considered statistically significant.

## ACKNOWLEDGMENTS

We would like to thank Dr. George Williams for inspiring our interest in nutrition modulation and ARDS. We would also like to thank Drs. Juli Bai, Holger Eltzschig, and Wenbo Li for their advice on the project direction.

