## Supplemental Figures 1-7 for "Ketogenic diet is protective during endotoxin-induced lung injury through the elevation of BHB"

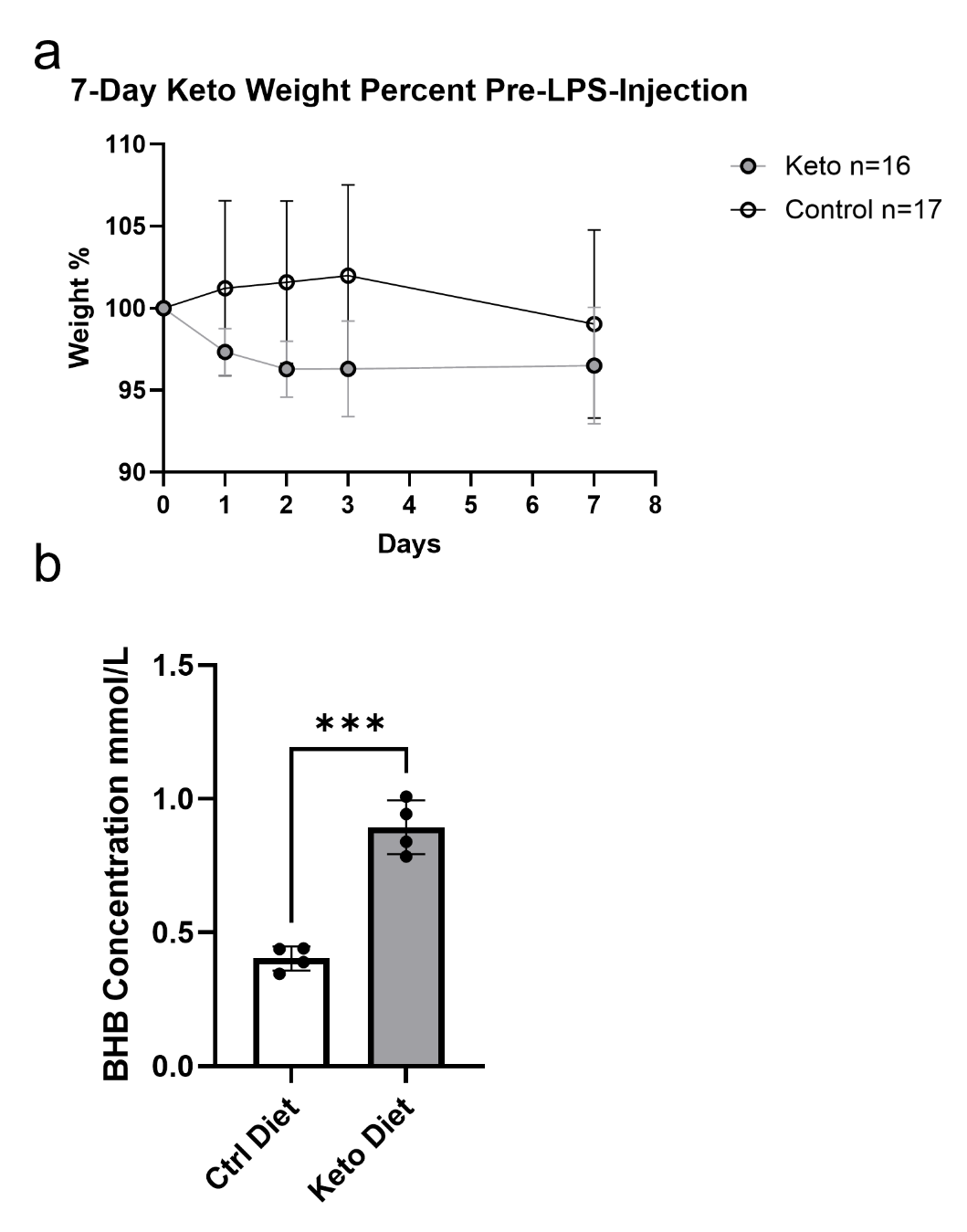

**Supplemental Figure 1: Phenotype of mice on ketogenic (keto) diet for 7 days prior to ARDS model. a.** 7-day weight percent curve of mice on a keto diet vs control mice before LPS-injection. (n=17 for the Control diet group, n=16 for the Keto diet group.) **b.** Compiled beta-hydroxybutyrate (BHB) concentration levels in control diet and keto diet mice blood serum samples from 2 separate trials. (n=4 per group, Unpaired t-test with Welch’s correction). All data are represented as mean ± SD; *P-value < 0.05

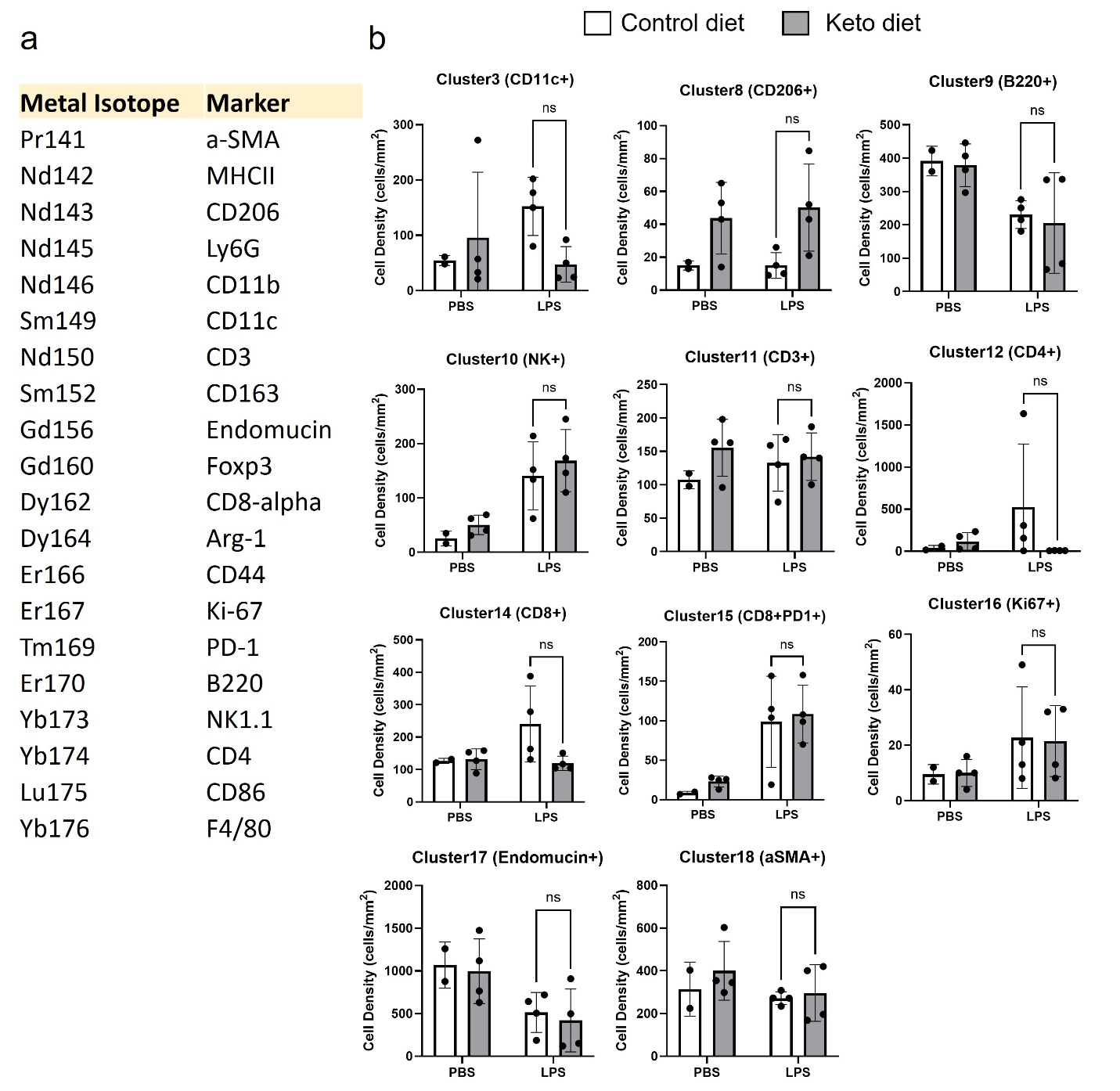

**Supplemental Figure 2: Additional Imaging Mass Cytometry (IMC) results. a.** Table of metal isotopes and their respective markers used for IMC. **b.** An additional list of significantly different immune cell populations based on the Uniform Manifold Approximation and Projection (UMAP) of mice on a 3-day LPS-induced ARDS, with four groups: Mice on a keto diet that were injected with or without LPS, and control diet mice that were injected with or without LPS (Figure 3). (n=2 in the Control diet PBS group, n=4 in the Ketogenic diet PBS group, n=4 in the Control diet LPS group, n=4 in the Ketogenic diet LPS group. Two-way ANOVA with Sidak test.). All data are represented as mean ± SD; ns P-value > .05.

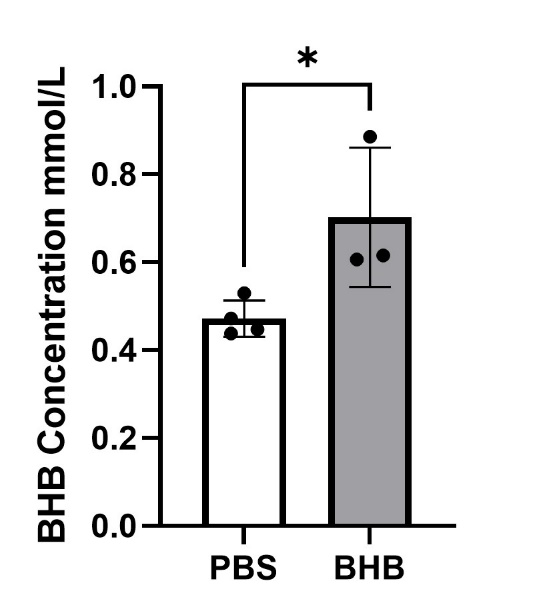

**Supplemental Figure 3: Confirmation of the efficacy of BHB treatment.** Compiled beta-hydroxybutyrate (BHB) concentration levels in PBS-injected and BHB-injected mice blood serum samples from 2 separate trials. (n=3-4 per group, Unpaired t-test with Welch’s correction). All data are represented as mean ± SD; *P-value < 0.05

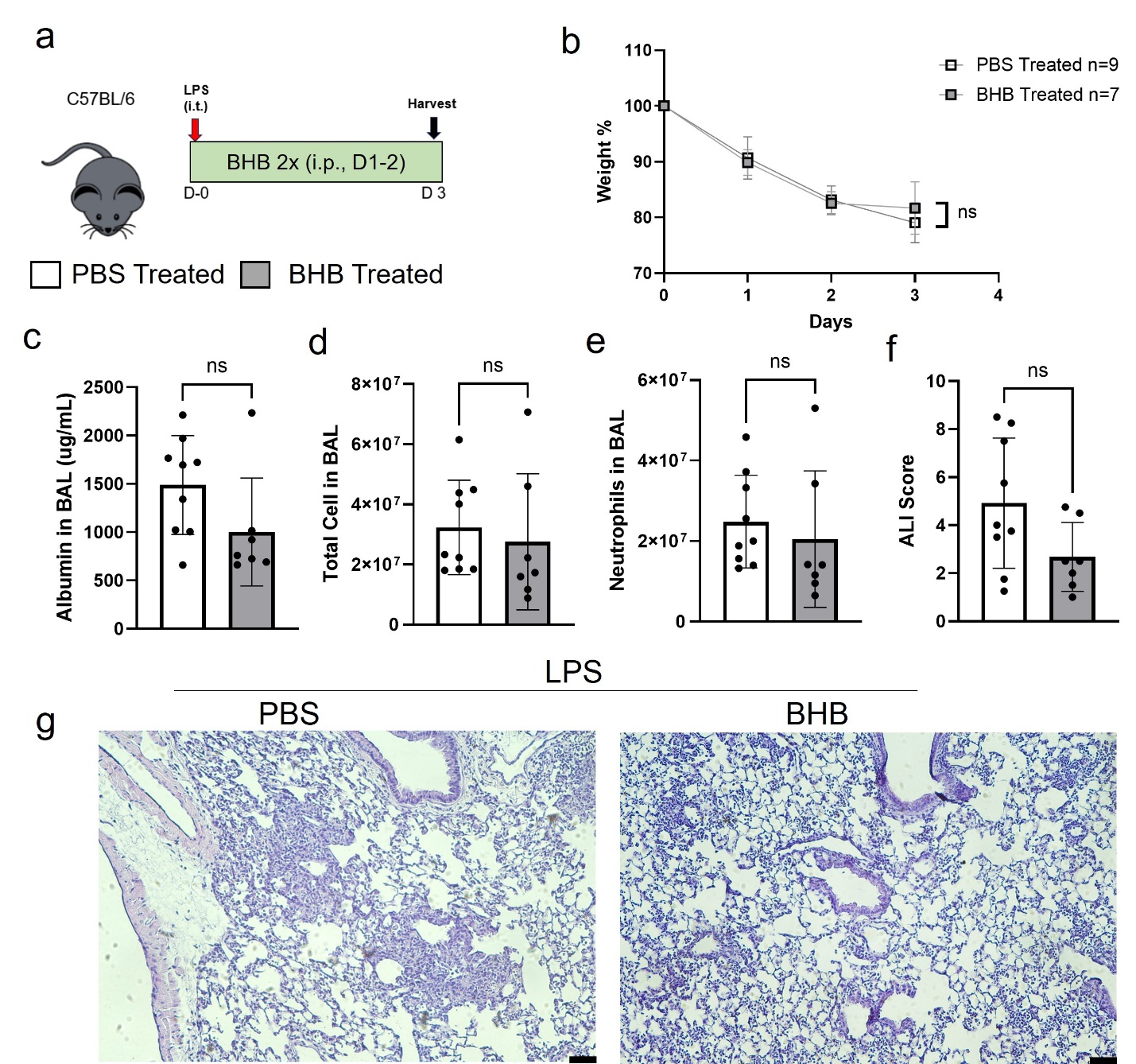

**Supplemental Figure 4: 2-day BHB injection phenotype. a.** An experimental layout. C57BL/6 mice had LPS instilled intratracheally on day 0 and followed for 3 days. Mice were injected with BHB twice a day for 2 days starting on day 1. Mice were harvested for their blood serum, BAL fluid, and lungs.  **b.** Weight loss curve of mice on a 3-day LPS-induced ARDS with two groups: Mice injected with BHB twice a day, and mice that were injected with PBS twice a day. (n=9 in the PBS group, n=7 in the BHB-treated group) **c.** Albumin levels extracted from BALF from mice on a 3-day LPS-induced ARDS. (n=7-9 per group, unpaired t-test with Welch’s correction.) **d.** Total cell count of BALF from mice on a 3-day LPS-induced ARDS. (n=7-9 per group, unpaired t-test with Welch’s correction.) **e.** Neutrophil cell count of BALF cytospin from mice on a 3-day LPS-induced ARDS. (n=7-9 per group, unpaired t-test with Welch’s correction.) **f.** ARDS score of the H&E-stained images. (n=7-9 per group, unpaired t-test with Welch’s correction.) **g.** 10x magnification of images of H&E staining of the lungs. **h.** Subcategories that were used for the ARDS scoring of the H&E-stained slides include: alveolar overdistension, atelectasis, cellularity, hemorrhage, perivascular edema, and peribranchial edema. (n=7-9 per group, unpaired t-test with Welch’s correction.) All data are represented as mean ± SD; *P-value < 0.05; ns p-value >0.05.

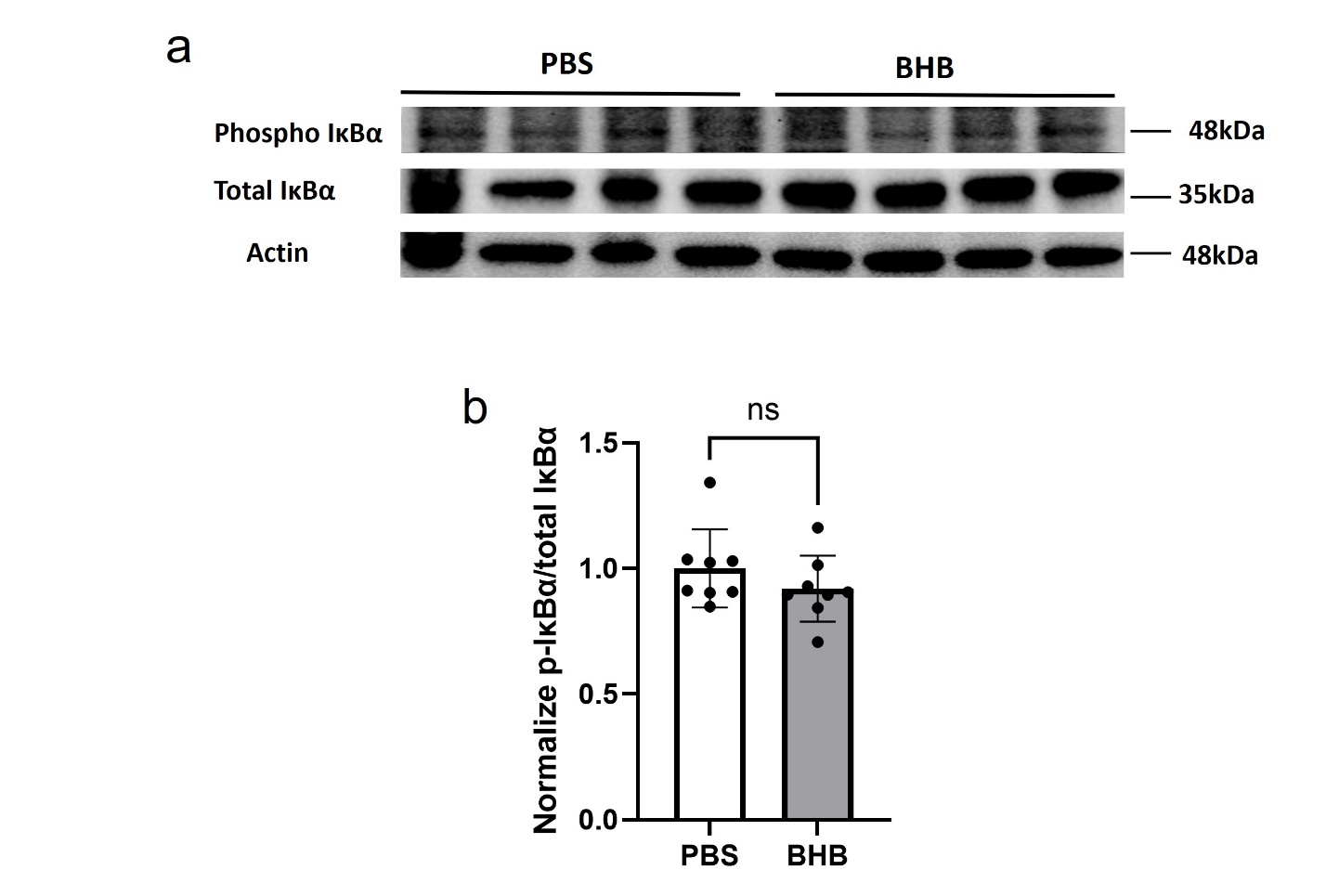
**Supplemental Figure 5: Phosphorylation of IκBα in lung tissue from BHB post-ARDS treatment.** C57BL/6 mice had LPS instilled intratracheally on day 0 and followed for 4 days. Mice were injected with BHB twice a day for 3 days starting on day 1. **a and b.** Image representative and densitometry quantification for Western blot for phosphorylated and total IκBα protein expression in lung tissue isolated from BHB-treated mice compared to PBS-treated mice after 4-day LPS-induced ARDS. Normalized phosphor-IκBα and total IκBα ratio was shown. (n=8 in the PBS-treated group, n=8 in the BHB-treated group. Unpaired t-test with Welch’s correction.) All data are represented as mean ± SD; ns p-value >0.05.

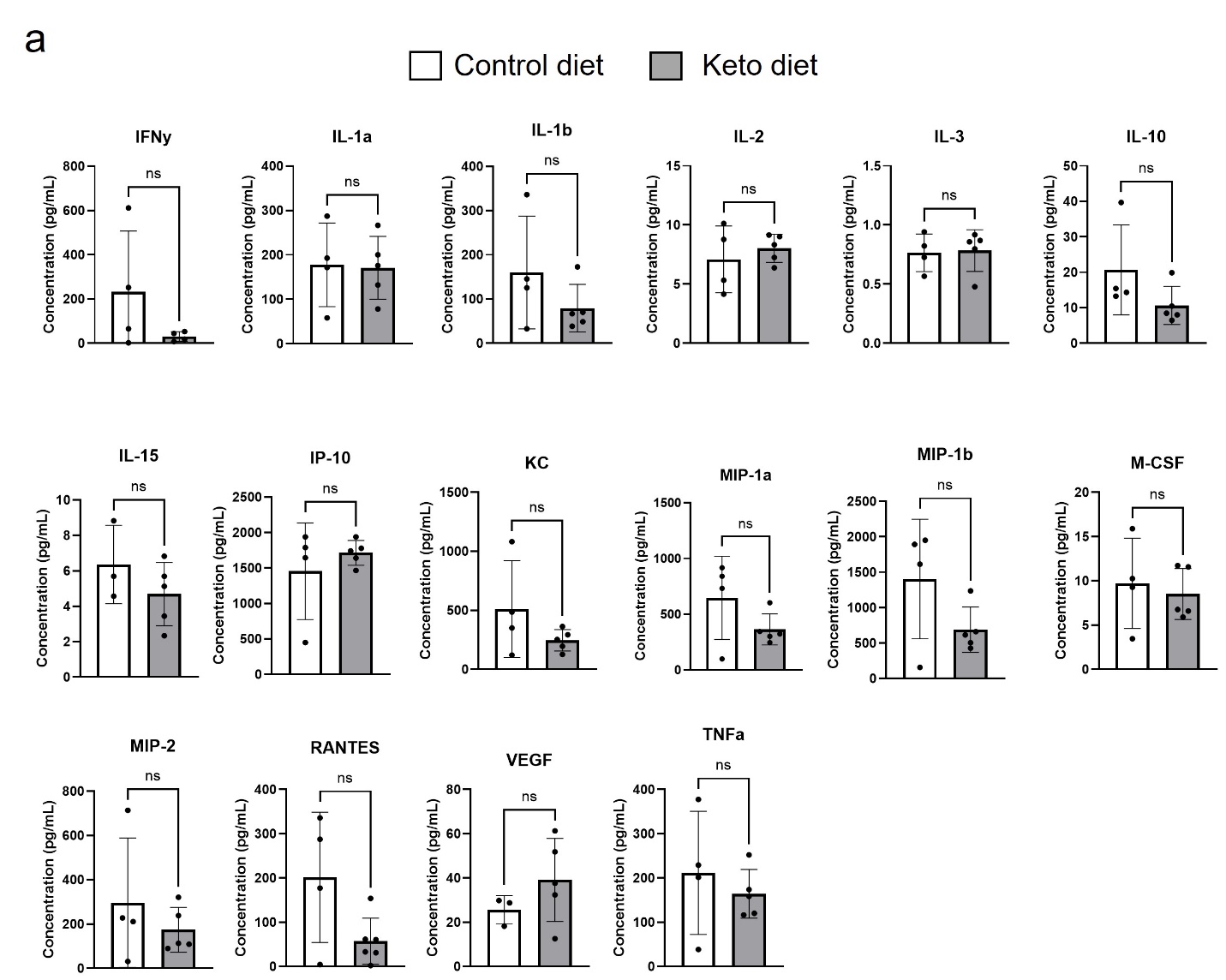

**Supplemental Figure 6: Additional Luminex results. a.** Additional list of different cytokines and chemokines from the Luminex mouse cytokine immunoassay in bronchoalveolar lavage fluid (BALF) from ketogenic (keto) diet mice compared to control mice after 3-day LPS-induced ARDS. (n=3-4 in the Control diet LPS group, n=5-6 in the Ketogenic diet LPS group. Unpaired t-test with Welch’s correction.) All data are represented as mean ± SD; *P-value < .05.

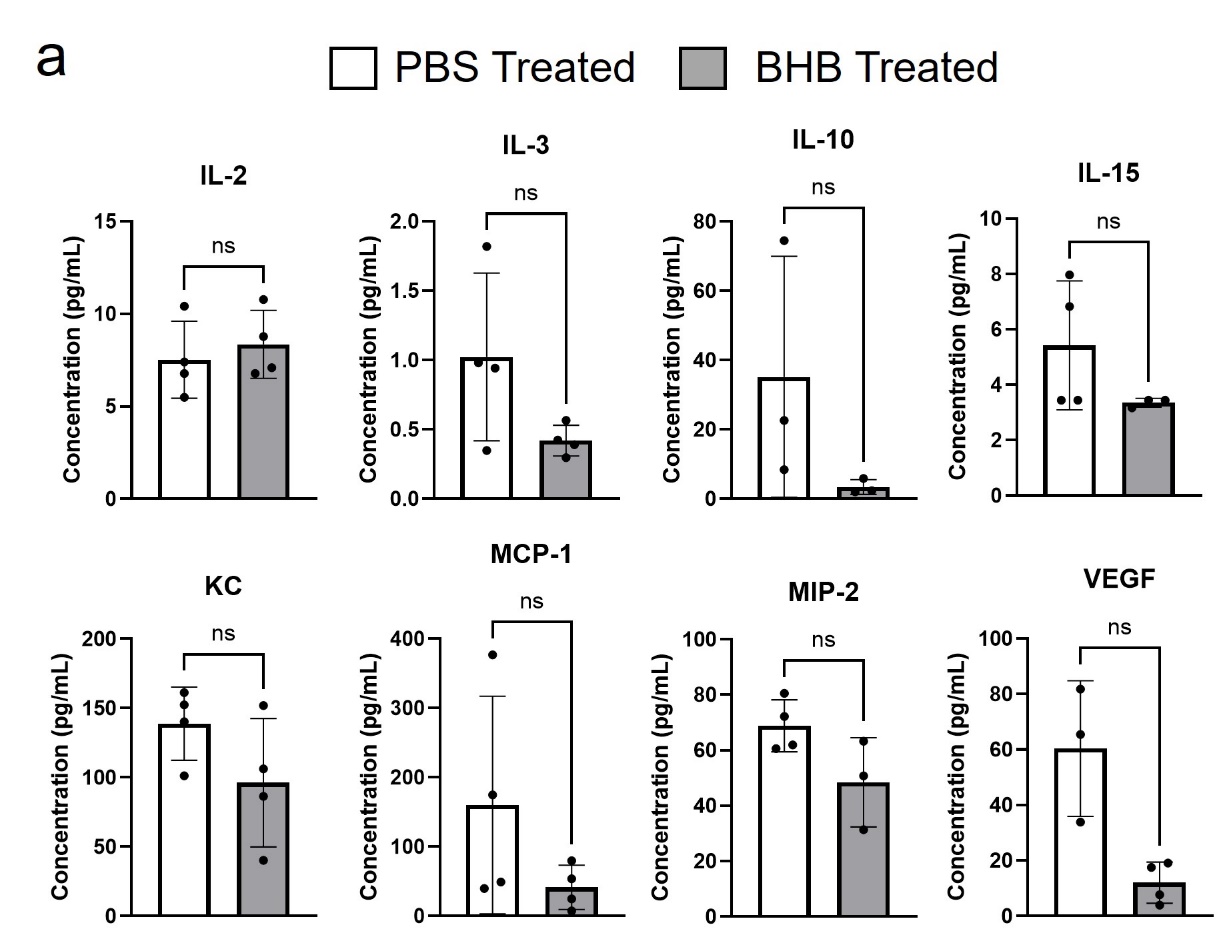

**Supplemental Figure 7: Additional Luminex results. a.** Additional List of different cytokines and chemokines from the Luminex mouse cytokine immunoassay in bronchoalveolar lavage fluid (BALF) from BHB-treated mice compared to PBS-treated mice after 4-day LPS-induced ARDS. (n=3-4 in the PBS-treated group, n=4 in the BHB-treated group. Unpaired t-test with Welch’s correction.) All data are represented as mean ± SD; *P-value < 0.05
